# Interval Timing is altered in male Nrxn1^+/-^ mice: A Model of Autism Spectrum Disorder

**DOI:** 10.1101/2025.08.14.670361

**Authors:** Kyle M. Roddick, Elias B. Habib, Richard E. Brown, Fuat Balcı

**Affiliations:** Department of Psychology, Mount Allison University, Sackville, NB, Canada; Department of Psychology and Neuroscience, Dalhousie University, Halifax, NS, Canada; Department of Epidemiology, Dalhousie University, Halifax, NS, Canada; Department of Biological Sciences, University of Manitoba, Winnipeg, MB, Canada

**Author notes:** Corresponding author Dr Kyle Roddick Department of Psychology Mount Allison University Sackville, NB, Canada.

**Keywords:** Interval Timing, Neurexin, Autism Spectrum Disorder, Mice, Genetic Murine Models

## Abstract

Autism spectrum disorder (ASD) is characterised by impaired social interactions and communication and increased repetitive and stereotypical behaviour. Neuroimaging shows functional abnormalities in brain areas involved in temporal processing of autistic individuals, and autistic individuals show deficits in interval timing. Neurexin (NRXN) mutations have been identified in a wide variety of neuropsychiatric disorders, including ASD, and Nrxn1^+/-^ mice possess a mutation that disrupts the α, β, and γ isoforms of Nrxn1, a gene involved in synapse structure. We investigated the interval timing abilities of the Nrxn1^+/-^ mouse model of ASD in the peak interval procedure using a 15-second target interval and compared their performance with that of Nrxn1^+/+^ and Nrxn1^ΔS5/-^ rescue mice. Two-month-old male Nrxn1^+/+^ (C57BL/6J), Nrxn1^+/-^, and Nrxn1^ΔS5/-^, mice were trained to obtain sucrose liquid rewards 15s after the onset of a discriminative stimulus (discrete fixed-interval training), and their timing responses were tested in non-reinforced probe trials. Our analysis of responses in individual trials revealed that Nrxn1^+/-^ mice had overall earlier timing responses. This difference was manifested as earlier termination of responding in terms of the response curves. These findings are consistent with leftward shifts observed with experimental animal models of ASD. In conclusion, we believe that these results are indicative of a biased long-term memory in the Nrxn1^+/-^ mouse model of ASD and may capture the timing deficit observed in autistic individuals.

**Lay Summary:** Neurexins help nerve cells connect and communicate with each other, and changes in these genes are often seen in people with autism. Mice with a change in their neurexin 1 gene, called Nrxn1^+/-^ mice, show autism-like behaviours. In a test that involves judging time, these mice respond early, similar to some people with autism. This study helps develop our understanding of how interval timing is affected in ASD.

## INTRODUCTION

Autism spectrum disorder (ASD) is a neurodevelopmental condition that is associated with a variety of deficits in social functions, cognition, sensory processing, and behavioral flexibility (American Psychiatric Association, 2022). In light of anecdotal evidence (Allman & DeLeon, 2009), it is possible that disrupted interval timing, namely the ability to keep track of time intervals in the seconds to minutes range, is at least partially responsible for ASD symptomatology, particularly in domains where temporal information is imperative (e.g., social cognition, communication, and predictive coding). Consequently, interval timing has become a topic of interest in the clinical domain (for review, see Allman & Meck, 2012; Casassus et al., 2019).

Allman et al. (2011) tested autistic children in a temporal bisection task that required participants to compare test durations as short or long based on their proximity to reference durations. Children were tested with two different sets of reference durations (1-4s and 2-8s). The psychometric functions of the autistic children were shifted leftward compared to the typically developing (TD) children for both reference duration sets. In addition, the psychometric functions were flatter in ASD children for the 2-8s reference durations (see Karaminis et al., 2016 for reduced timing sensitivity in ASD). In another set of tasks, where participants were asked to reproduce target intervals, Szelag et al. (2004) found that the time reproductions of ASD children differed more from the target durations than TD children (see Brenner et al., 2015; Maister & Plaisted-Grant, 2011 for disrupted temporal accuracy). Despite the findings that point to poor timing ability in autistic individuals, a number of studies failed to find any difference.

For instance, Gil et al. (2012) did not find any difference between ASD and TD children who were tested in the temporal bisection task with a wide range of reference interval pairs (also see Jones et al., 2017 for lack of differences in a temporal bisection in an adult sample). In addition, neither Wallace and Happé (2008) nor Doenyas et al. (2019) found any disruption in the timing performance of ASD children using a wider range of tasks (e.g., temporal reproduction, rhythmic timing). Overall, the results gathered from human participants are inconsistent regarding whether or not interval timing is affected in ASD (for a systematic review of studies on time perception in ASD individuals, see Casassus et al., 2019).

Few studies have investigated how interval timing is affected in animal models of ASD based on prenatal exposure to valproic acid (see Patterson, 2011). In one study, male and female CF1 mice were prenatally exposed to valproic acid (VPA) and later tested using the dual peak procedure with 15-second and 45-second target intervals (Acosta et al., 2018). In the peak interval procedure, mice were trained to anticipate reward delivery contingent upon their first response after a fixed delay following the onset of a discriminative stimulus. In peak interval trials, the discriminative stimulus is presented for a longer duration; the reward is omitted, and the modulation of anticipatory responding as a function of stimulus duration is examined. The response rate is typically maximal at the time of reward availability. In the dual peak interval procedure used by Acosta et al. (2018), two different levers were associated with different delays until reinforcement: 15 seconds and 45 seconds. They found that both male and female mice that were prenatally exposed to VPA (and weaned with other VPA mice) had leftward shifts in their timed responses; female VPA mice had a leftward shift for the 15s interval only, while male VPA mice had a leftward shift in both 15s and 45s intervals. In addition, both male and female VPA mice had lower temporal precision and lower peak amplitude than control mice. DeCoteau and Fox (2021) tested male Wistar rats prenatally exposed to VPA on fixed-interval (FI) temporal bisection, peak interval, and intertemporal choice timing tasks. They found a leftward shift in the middle times of the VPA rats on the peak interval task, and a trend towards a leftward shift in the FI temporal bisection task. In summary, these studies demonstrate a clear timing deficit in animal VPA models of ASD.

An alternative approach that offers higher construct validity than VPA exposure is the use of genetically modified animals that replicate the genetic mutations associated with the human condition. To this end, mutations to neurexin genes that control the production of presynaptic cell adhesion molecules may provide a better model of ASD. The neurexins play a critical role in synaptic development (Krueger et al., 2012), and are associated with a number of neuropsychiatric conditions, including ASD and schizophrenia (Gauthier et al., 2011; Gomez et al., 2021; Hu et al., 2019; Kasem et al., 2018; Reissner et al., 2013). Neurexin 1 is a molecule that acts to recruit or stabilise neurotransmitter receptors (Craig & Kang, 2007). NRXN1 point mutations have been identified in ASD patients (Onay et al., 2016) and NRXN1 exon deletions are estimated to increase the risk of ASD by approximately 20-fold (Dabell et al., 2013). Several previous studies have examined the effect of Nrxn1α knockout in mice, and some have reported ASD-like phenotypes. For instance, Etherton et al. (2009) found impaired prepulse inhibition and altered nest-building behavior, as well as increased self-grooming, in Nrxn1α knockout mice.

However, they did not observe differences in other tasks, including social behavior and anxiety. Grayton et al. (2013) found altered social behavior in male and female homozygous (but not hemizygous) Nrxn1α knockout mice, which displayed altered social approach and reduced social investigation, and males also showed increased aggressive behavior. Dachtler et al. (2015) also found decreased social behavior in heterozygous Nrxn1α knockout mice, which showed no preference for a novel mouse in a social approach task. Armstrong et al. (2020) examined Nrxn1α knockout mice across different ages and found that homozygous Nrxn1α knockout mouse pups produced shorter ultrasonic vocalizations than their hemizygous and wild-type littermates. Abnormal social behaviour, such as decreased investigative and affiliative behaviours and increased levels of aggression were observed in male homozygous and hemizygous Nrxn1α knockout mice, both as juveniles and as adults.

These studies used knockouts affecting only Nrxn1α, leaving the β and ɣ isoforms of Nrxn1 unaffected. Lu et al. (2025) tested mice hemizygous for a knockout of Nrxn1α, β, and ɣ and found deficits in excitatory synaptic transmission in the hippocampus, lower protein levels of Nrxn1, and typical ASD-like behavioural deficits, including increased rearing and grooming, and decreased nest quality, but no deficits in social behaviour. Lu et al. (2023) had shown that exclusion of splice site 5 of Nrxn1 boosts hippocampal excitatory synapse density and transmission and resulted in behavioural changes opposite to those expected in ASD models, including increased pup ultrasonic vocalizations and decreased grooming . The use of splice site 5 exclusion as a therapeutic approach to the effects of hemizygous knockout of Nrxn1 alleviated the deficits in excitatory synaptic transmission and the behavioural deficits, and partially alleviated the reduced Nrxn protein levels (Lu et al., 2025).

To our knowledge, no study has investigated how interval timing, a potential biomarker of ASD, is affected in a genetic mouse model of ASD, and certainly not in a rescue model. The current study bridges this gap by investigating the timing behavior of a novel mouse model of ASD, which is hemizygous for the α, β, and γ isoforms of neurexin 1, as well as genetic rescue, Nrxn1^ΔS5/-^, and WT mice on the peak interval timing task. We hypothesized that the Nrxn1^+/-^ mice would have abnormal timing performance compared to the Nrxn1^+/+^ mice, and that the Nrxn1^ΔS5/-^ mice would be similar to the Nrxn1^+/+^ mice.

## METHODS

### Subjects

Forty-eight male mice, 16 C57BL/6J (Nrxn1^+/+^), 16 Nrxn1^+/-^ mice, and 16 Nrxn1^ΔS5/-^ mice, were tested at 2 to 3 months of age. The Nrxn1^+/-^ mice are hemizygous for a 140-bp deletion that knocks out the α, β, and γ isoforms of Nrxn1, while the Nrxn1^ΔS5/-^ mice, a proposed genetic rescue of the Nrxn1^+/-^ mice (Lu et al., 2025), have the same 140-bp deletion to one copy of Nrxn1, and the remaining copy has an exclusion of splice site 5, shown to upregulate Nrxn1 protein levels and alleviate synaptic transmission and behavioural deficits caused by the 140-bp deletion (Lu et al., 2023, 2025). Nrxn1^+/+^ and Nrxn1^+/-^ mice were bred by mating wildtype Nrxn1^+/+^ females and transgenic Nrxn1^+/-^ males, while Nrxn1^ΔS5/-^ mice were bred by mating Nrxn1^ΔS5/-^ and Nrxn1^ΔS5/ΔS5^ mice. All mice were bred in-house at Dalhousie University from mice obtained from Dr. Ann Marie Craig at the University of British Columbia. Genotypes were determined using PCR with DNA from ear punches by Dr. Chris Sinal at Dalhousie University. The mice were weaned at 30 days of age and separated into same-sex groups of 2 to 4 siblings, housed in 30 cm × 18 cm × 12 cm polycarbonate cages with wire tops, and had ad lib access to food (Purina Rodent Laboratory Chow #5001) and water. The cages had wood chip bedding, a 5 cm diameter by 7 cm long PVC tube for enrichment, and paper strips were provided as nesting material. The colony room was maintained at 20 ± 2°C on a reversed 12-hour light/12-hour dark cycle, with lights off from 9:30 am to 9:30 pm. All testing was performed during the dark phase of the cycle. All procedures were approved by the Dalhousie University Committee on Laboratory Animals and were conducted in accordance with the guidelines of the Canadian Council on Animal Care (protocol #18-096).

### Apparatus

The apparatus and procedures were the same as those described by Gür, Fertan, Kosel et al. (2019). Testing was conducted in a mouse nine-hole box (Cambridge Cognition Ltd., England) with a lick tube attached to a peristaltic pump, house lights, and speakers (mounted on both sides of the inner walls). The box was placed in a sound- and light-attenuating chamber with a video camera on top, allowing the behaviour of the mice to be observed without disrupting the test procedure. The test chamber contained a grid floor with a removable tray underneath. Six of nine holes (Holes 1–3, 7–9) were plugged, and the remaining three holes in the middle (Holes 4–6) were left open. Each of the three holes and the reinforcement tube/tray could be illuminated, and nose pokes to the open holes were detected via infrared beams. Computer software (Cambridge Cognition Ltd., England) was used to control the test box and record time-stamped nose pokes.

### Procedure

#### Water restriction

Two days before the first training session started, the mice were separated into individual cages, and their water intake was restricted while maintaining each animal at 85% of its ad libitum weight. On the first day of water deprivation, the water bottles were removed from the cages, and each mouse was given powdered rodent chow that was mixed with tap water (mash). The weight of the mice was maintained at the desired level by providing mash (on average, 2.5 mL) after each test. Subjects also received a 5% sucrose solution as a reward during testing. During water restriction, subjects had *ad lib* access to food in their home cages.

#### Peak Interval Procedure Magazine training

Mice received one 20-minute session of magazine training per day for two consecutive days. At the start of the sessions 0.025 ml of 5% sucrose water was delivered via the peristaltic pump, followed by 0.02 ml every 40 s during the session (note that 0.7 ml indicated in Gür, Fertan, Alkins, et al., 2019; Gür, Fertan, Kosel, et al., 2019 was the approximate total amount of water received during magazine training and not the amount delivered every 40 s). The reward tray was illuminated throughout the sessions.

#### Fixed interval training

Mice received one 20-minute session of fixed interval (FI) training per day for four consecutive days. In each trial, reinforcement (0.03 ml of 5% sucrose water) was delivered contingent upon the first nose poke into the central nose poke hole after 15 s since the onset of the discriminative stimulus (i.e., the onset of the light in the central nose poke hole). Trials with no response after the fixed interval were terminated after 45 s. Following a nose poke response, the light was extinguished and an inter-trial interval (ITI) of 20 s fixed ± a uniformly distributed random variable with a mean of 10 s commenced.

#### Peak interval testing

Mice received one 20-minute session of peak interval (PI) testing per day for 30 consecutive days. PI trials were randomly mixed with FI trials with a 1:2 ratio of PI:FI trials. The PI trials began with the illumination of the central nose poke hole and lasted 45 seconds. At the end of the PI trial, the light in the central nose poke hole was turned off, and an ITI started. No reinforcement was given during PI trials. Due to a one-day interruption in the testing of one batch of mice (5 Nrxn1^+/-^ mice and 2 Nrxn1^+/+^ mice), the data collected from these mice the day after the interruption were excluded from the dataset prior to data analysis.

#### Data Analysis

Data analysis was conducted with MATLAB (version 25.1 (R2025a); The MathWorks Inc., 2025) and R (version 4.5.1; R Core Team, 2025).

#### Analysis of the individual trial data

In individual PI trials, subjects exhibit a response pattern that differs from the smooth bell-shaped average response curves (Church et al., 1994). In each trial, responding in the steady state occurs as a pattern composed of three stages. At the beginning of the trial, the response rate is low, and as the time of reinforcement availability approaches, it increases abruptly (start time).

Since reinforcement is omitted in the PI trials, sometime after the time that the reinforcement should be available, the response rate abruptly declines (stop time). The period of high response rates is typically clustered around the time of reinforcement availability, indicating the period when there is a high expectancy of reinforcement delivery on that trial. The interval between the stop and start times is referred to as the spread, which reflects timing uncertainty. The average of the start and stop times reflects the targeted interval in that trial.

The primary aim of the individual trial analysis was to estimate the trial time at which animals shift from break to a run (start time) and the run back to a break (stop time). In other words, the time when the subject starts anticipating and stops anticipating the reward delivery (following its omission), respectively. The period before the run, the first break period prior to the subject initiating its high rate of anticipatory responding, should have lower-than-average response rates. The run should have higher-than-average response rates, and the second break period, following the subject terminating its high rate of anticipatory responding, should again show lower-than-average response rates. To detect the start and stop times, we used a variant of the algorithm presented in Church, Meck, and Gibbon (1994), which assumed a second start time.

Additionally, the coefficients of variation for the start, stop, and spread times were compared. Comparisons between genotypes for these measures were made using the last five sessions of the trial, performed with one-way ANOVA.

As recommended by Karson & Balcı (2021), based on Gibbon & Church’s (1990) theoretical work, the correlation coefficients between the start and stop times, the start times and the spreads, and the middle times and spreads were calculated for each mouse using the data from the last five sessions. Comparisons between genotypes for each of these three correlation coefficients were performed using one-way ANOVA.

#### Analysis of the peak response curve

The average response curves in the PI procedure are nearly bell-shaped, with the peak located around the 15s delay of reinforcement availability. The latency of the peak (i.e., peak time) reflects the time of maximum expectancy for the reinforcement delivery (timing accuracy), the width of the response curve (i.e., spread) measures timing variability, and the amplitude of the response gradient reflects the motivation level of the subjects, independent of the timing performance (Balcı, 2014; Roberts, 1981). For parameter estimations from the average response curve or gradient, we use a method previously applied by Balcı et al. (2009). The peak response curve was determined using the trials from the final five sessions. Initially, the average response rate data of each mouse was expressed in 1-s bins. The overall response rate was calculated for each mouse by taking the mean of the binned average response rates that formed the average response curve. The amplitude was also determined from the non-normalized average response curve as the height of the curve at its peak. For the estimation of the peak time and response curve width, on the other hand, each response curve was normalized by its amplitude for each subject then smoothed by a window of three bins, which replaces the average value in 1-s bin with the average of the three bins surrounding it (except for the endpoints of the data). The smoothing method using a moving average is applied to reduce noise in the individual response curves while preserving their shape. The global maximum of the normalized response curve is the peak time. The first point that exceeded the normalized response rate of 0.70 is determined as the start point. If there is a decrease below the normalized rate of 0.50 between this point and the peak time, the start point is determined after this point. The first point at which the normalized response rate falls below 0.70 after reaching its peak is determined as the stop point. The width of the response curve was determined as the difference between the average stop and start points derived from the response curve. Each parameter was compared between genotypes using one-way ANOVA.

#### Analysis of response rates during ITI and the discriminative stimulus

The mean number of responses made by the mice across all sessions was compared. To evaluate whether the mice learned to associate the discriminative stimulus with the reinforcement, the ratio of nose pokes during the presentations of the discriminative stimulus (light) to the number of nose pokes during the ITIs was calculated. A higher ratio is indicative of better associative learning (e.g., Papachristos & Gallistel, 2006); however, this measure is confounded by timing imprecision (an aspect of poor timing performance). Finally, the overall locomotor activity was quantified as the number of infrared beam breaks placed in front of the choice wall. One-way ANOVA was used to examine genotype differences in these metrics.

## RESULTS

### Well-Timed Trials

During the final block of testing, there was no significant genotype difference in terms of the number of poorly timed trials (i.e., start times > 15s or stop times < 15s; *F*_(2,45)_ = 1.34, *p* = 0.273, 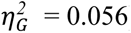). Thus, all trials were included in the subsequent analyses.

### Single Trial Analysis

The single-trial parameters were analyzed using one-way ANOVA, followed by planned comparisons between Nrxn1^+/+^ and Nrxn1^+/-^, and between Nrxn1^+/-^ and Nrxn1^ΔS5/-^ mice.

### Start and Stop Times

While there was no main effect of genotype on start times (*F*_(2,45)_ = 2.68, *p* = 0.079, 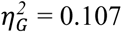; Figure 1A), the planned comparisons showed that the Nrxn1^+/-^ mice (8.25 ± 2.03) had an earlier start time than the Nrxn1^ΔS5/-^ mice (10.2 ± 2.85; *t*_(45)_ = -2.22, *p* = 0.0315, *d* = -0.782). There was a main effect of genotype on stop times (*F*_(2,45)_ = 4.26, *p* = 0.02, 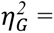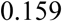; Figure 1C), with the Nrxn1^+/-^ mice (25.9 ± 2.24) having earlier stop times than the Nrxn1^ΔS5/-^ mice (28.6 ± 2.78; *t*_(45)_ = -2.92, *p* = 0.00548, *d* = 1.05).

**Figure 1.**
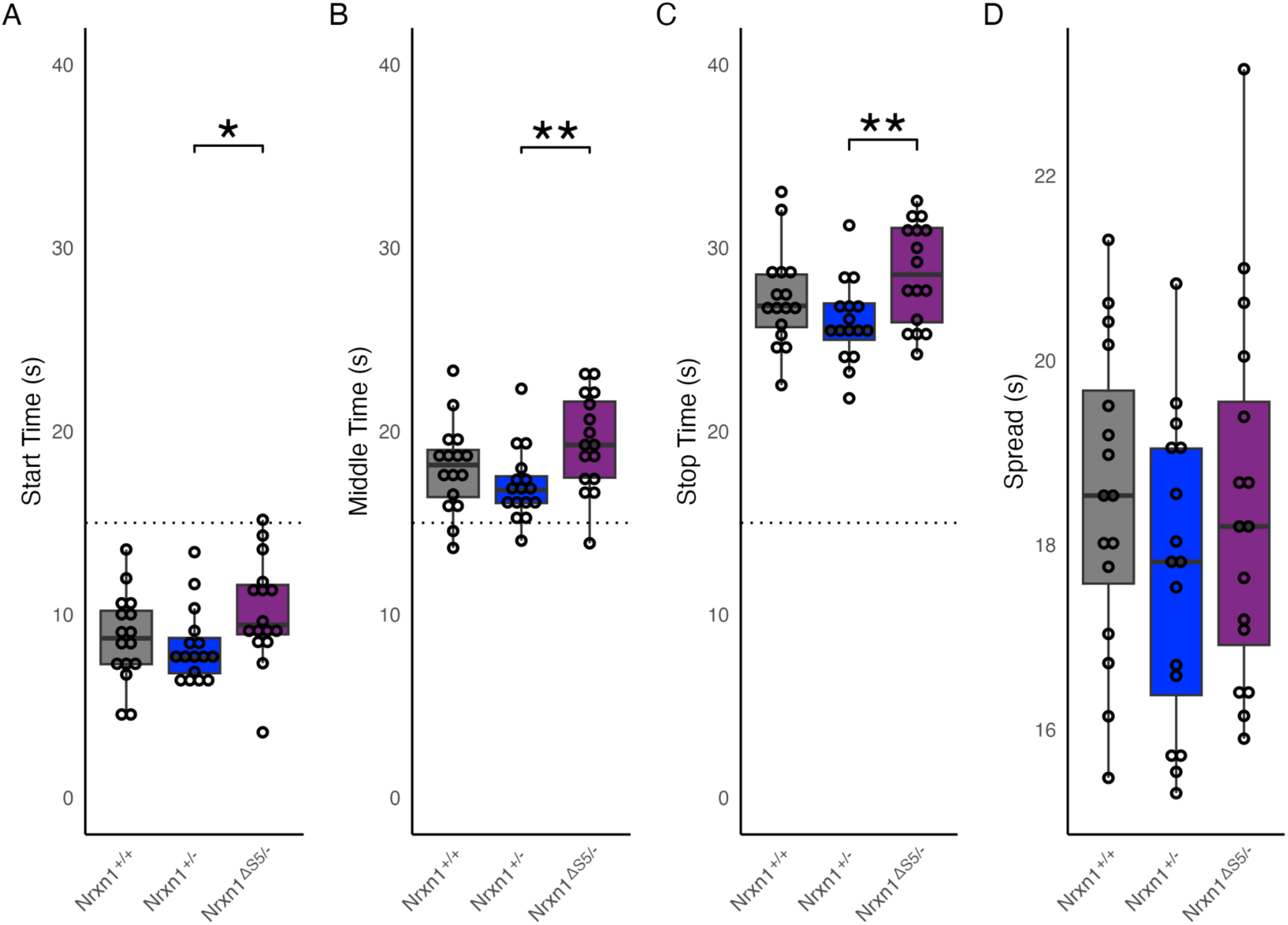
The comparison of single-trial analysis between the Nrxn1^+/+^, Nrxn1^+/-^, and Nrxn1^ΔS5/-^ mice for A) start times, B) middle times (i.e., [start time + stop time] / 2), C) stop times and D) spread (i.e., stop time - start time). The visual inspection of these values show an overall leftward shift in the timing responses of Nrxn1^+/-^ mice particularly compared to Nrxn1^ΔS5/-^ mice. These shifts were not accompanied by wider response periods that would result from higher timing uncertainty. The boxes indicate the interquartile ranges, with the medians indicated by the horizontal lines, whiskers show the range of data points out to a maximum of 1.5 interquartile ranges, and the individual data points are plotted over the boxes (0.05 * 0.01 ** 0.001 *** 0).

In order to interpret the timing precision at a more granular level, we analyzed the CV of start and stop times. There was an effect of genotype on the coefficient of variation (CV) of the start times (*F*_(2,45)_ = 4.09, *p* = 0.023, 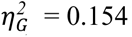), with the Nrxn1^+/+^ mice (0.802 ± 0.117) having a lower CV values than the Nrxn1^+/-^ mice (0.956 ± 0.214; *t*_(45)_ = -2.57, *p* = 0.0136, *d* = -0.894), and the Nrxn1^+/-^ mice (0.956 ± 0.214) having greater CV values than the Nrxn1^ΔS5/-^ mice (0.814 ± 0.164; *t*_(45)_ = 2.37, *p* = 0.022, *d* = 0.747). There was no significant effect of genotype on the CV of stop times (*F*_(2,45)_ = 0.865, *p* = 0.428, 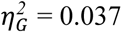; Nrxn1^+/+^: 0.271 ± 0.0743; Nrxn1^+/-^: 0.28 ± 0.0585; Nrxn1^ΔS5/-^: 0.298 ± 0.0367).

### Middle & Spread Times

There was an effect of genotype on middle times (*F*_(2,45)_ = 3.86, *p* = 0.028, 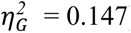), with the the Nrxn1^+/-^ mice (17.1 ± 1.97) having earlier middle times than the Nrxn1^ΔS5/-^ mice (19.4 ± 2.63; *t*_(45)_ = -2.76, *p* = 0.008, *d* = 0.989; Figure 1B). There was no effect of genotype on spread (*F*_(2,45)_ = 1.03, *p* = 0.366, 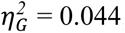; Figure 1D).

### Correlational Analysis

Correlational analyses are informative in terms of the sources of variability in timing behavior. Pearson’s correlations between the start and stop times, the start time and the spread, and the spread and the middle time were also examined. Genotype differences in correlation coefficients were analyzed with ANOVAs. Start-stop time correlations (Nrxn1^+/+^: *r*_(629)_ = 0.7, *p* < 0.0001; Nrxn1^+/-^: *r*_(611)_ = 0.72, *p* < 0.0001; Nrxn1^ΔS5/-^: *r*_(576)_ = 0.69, *p* < 0.0001) and start time-spread correlations (Nrxn1^+/+^: *r*_(629)_ = -0.31, *p* < 0.0001; Nrxn1^+/-^: *r*_(611)_ = -0.43, *p* < 0.0001; Nrxn1^ΔS5/-^: *r*_(576)_ = -0.32, *p* < 0.0001) were significant for all genotypes, and different genotypes of mice did not differ on either start-stop time correlations (*F*_(2,45)_ = 1.26, *p* = 0.293, 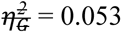), or start- spread correlations (*F*_(2,45)_ = 2.21, *p* = 0.122, 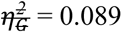). While the middle-spread correlations were significant for the Nrxn1^+/+^ mice (*r*_(629)_ = 0.1, *p* = 0.0106) and the Nrxn1^ΔS5/-^ mice (*r*_(576)_ = 0.094, *p* = 0.0234), they were not significant for the Nrxn1^+/-^ mice (*r*_(611)_ = -0.079, *p* = 0.052), resulting in a significant difference in the middle-spread correlations between genotypes (*F*_(2,45)_ = 3.27, *p* = 0.047, = 0.127), with the Nrxn1^+/-^ (-0.0996 ± 0.259) mice showing lower correlations than both the Nrxn1^+/+^ mice (0.0909 ± 0.224; *t*_(45)_ = 2.23, *p* = 0.0309, *d* = -0.787) and the Nrxn1^ΔS5/-^ mice (0.0885 ± 0.242; *t*_(45)_ = -2.2, *p* = 0.033, *d* = -0.751).

### Response Curve Analysis

The parameters calculated from the average response curves (Figure 2A) were compared between genotypes using ANOVAs. There was an effect of genotype on stop times (*F*_(2,45)_ = 3.26, *p* = 0.048, 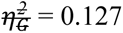; Figure 2E), with planned comparisons showing the Nrxn1^+/-^ mice (24.3 ± 3.66) had earlier peaks than the Nrxn1^ΔS5/-^ mice (30.2 ± 8.13; *t*_(45)_ = -2.47, *p* = 0.0172, *d* = -0.932). There was also an effect of genotype on the amplitude at the tail of the response curve (i.e., 30s; *F*_(2,45)_ = 3.37, *p* = 0.043, 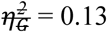; Figure 2G), with planned comparisons showing the Nrxn1^+/+^ mice (0.599 ± 0.249) had a greater amplitude than the Nrxn1^+/-^ mice (0.407 ± 0.259; *t*_(45)_ = 2.18, *p* = 0.0342, *d* = 0.757), as well as the Nrxn1^ΔS5/-^ mice (0.61 ± 0.239) having a greater amplitude than the Nrxn1^+/-^ mice (*t*_(45)_ = -2.31, *p* = 0.0258, *d* = 0.815). No other average response curve measures showed a significant difference between genotypes (*p*s ≥ 0.202).

**Figure 2.**
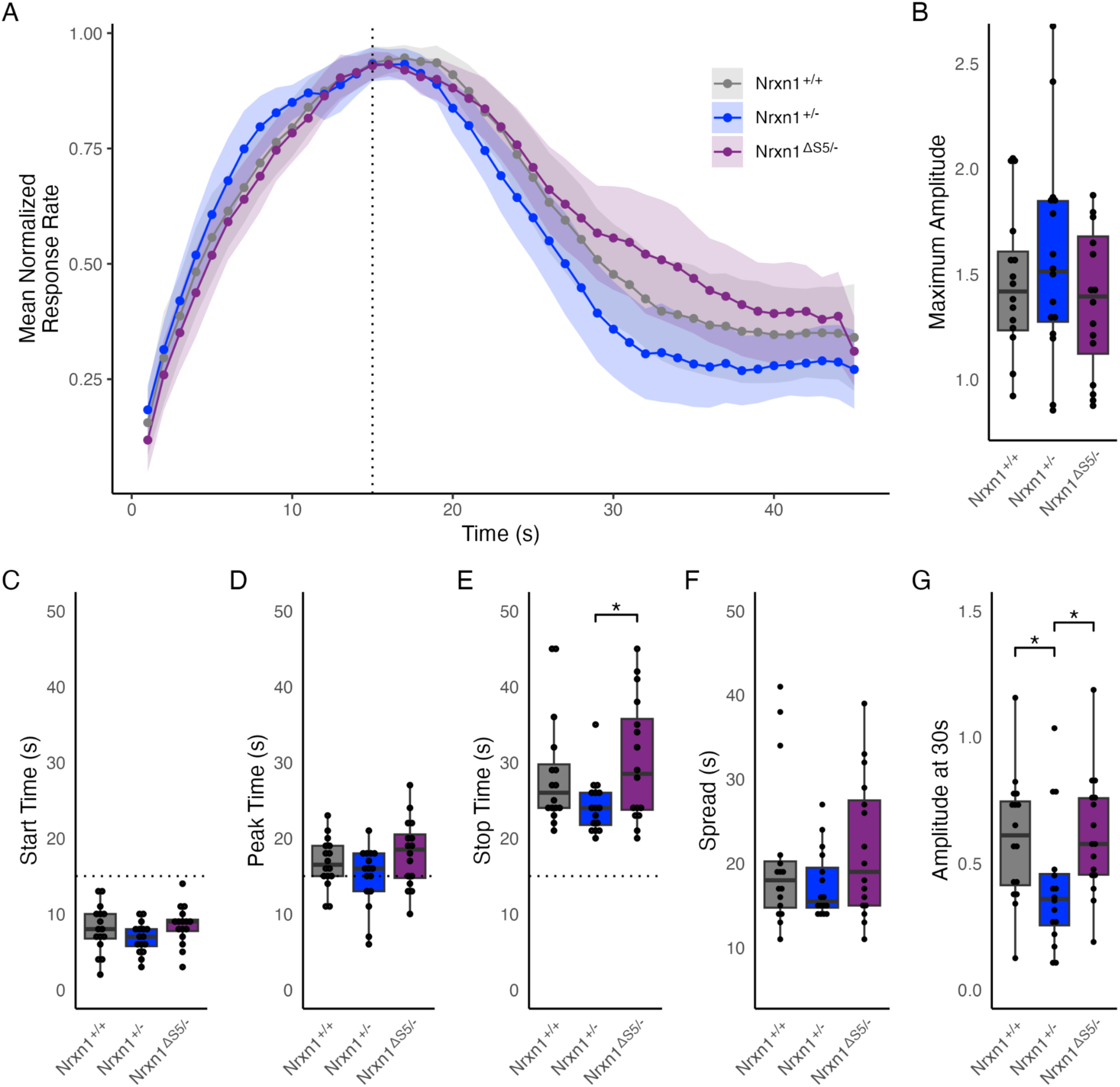
A) Average peak response curves of the Nrxn1^+/+^, Nrxn1^+/-^, and Nrxn1^ΔS5/-^ mice. The timing indices estimated from the average peak response curves are shown in the other panels for the same groups: B) maximum amplitude of the non-normalized response rate of the average response curve, C) start times, D) peak times, E) stop times, F) spread (i.e., stop times - start times) and G) response rate around the right-hand tail of the average response curve (i.e., 30s). For subplots B through G, the boxes indicate the interquartile ranges, with the medians indicated by the horizontal lines, whiskers show the range of data points out to a maximum of 1.5 interquartile ranges, and the individual data points are plotted over the boxes (0.05 * 0.01 ** 0.001 *** 0).

### Response rates and activity throughout the session

Response rates and activity around the choice wall (measured by the total number of a single infrared beam placed before the choice wall) were analyzed with ANOVAs. During the final block of testing, there were no genotypes differences in the number of total nose pokes per session (Nrxn1^+/+^: 6381 ± 2106, Nrxn1^+/-^: 7111 ± 2225, and Nrxn1^ΔS5/-^: 6426 ± 2699; *F*_(2,45)_ = 0.481, *p* = 0.621, 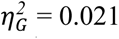). This was true when examining both nose pokes made while the discriminative stimulus was on (Nrxn1^+/+^: 4241 ± 1407, Nrxn1^+/-^: 4726 ± 1488, and Nrxn1^ΔS5/-^: 4245 ± 1759; *F*_(2,45)_ = 0.512, *p* = 0.603, 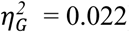), and during the inter-trial intervals (Nrxn1^+/+^: 2140 ± 700, Nrxn1^+/-^: 2385 ± 738, and Nrxn1^ΔS5/-^: 2181 ± 941; *F*_(2,45)_ = 0.430, *p* = 0.653, 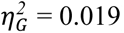). The ratio of the number of nose pokes made during the discriminative stimulus to the number of nose pokes during the ITIs did not differ (Nrxn1^+/+^: 1.98 ± 0.049, Nrxn1^+/-^: 1.98 ± 0.043, and Nrxn1^ΔS5/-^: 1.95 ± 0.033; *F*_(2,45)_ = 1.76, *p* = 0.183, 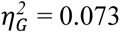). The total amount of activity around the choice wall did not differ (Nrxn1^+/+^: 4434 ± 1377, Nrxn1^+/-^: 4220 ± 1312, and Nrxn1^ΔS5/-^: 3822 ± 530; *F*_(2,45)_ = 1.19, *p* = 0.315, 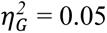), indicating similar levels of movement by each genotype in the testing apparatus.

## DISCUSSION

The current study investigated how timing behavior is altered in male Nrxn1^+/-^ mice, a novel genetic model of ASD, and compared their performance with that of WT and rescue mice Nrxn1^ΔS5/-^. Consistent with the findings of Allman et al. (2011), Acosta et al. (2018), and DeCoteau and Fox (2021), we observed that male Nrxn1^+/-^ mice exhibited a leftward shift in their timing behavior but interestingly, this difference was significant only with respect to the Nrxn1^ΔS5/-^ mice.

Acosta et al. (2018) found that female mice exhibited this effect only for the 15s interval, both in terms of the response curve and single-trial analysis. Male VPA mice had leftward shifts for both the 15s and 45s intervals, observed only in single-trial analysis. These shifts were found to be primarily due to changed start times, which suggests the motivational mediation of the effect (see Balcı, 2014). We observed a leftward shift in the middle times, which was accompanied by leftward shifts in both start and stop times (suggesting a true timing phenotype). In contrast, in the response curve analysis, the most prominent shift was in the stop times, as in Acosta et al. (2018). Thus, despite minor differences overall, the timing behavior (e.g., peak times) is altered in the same direction in both mouse models of ASD.

Finally, Acosta et al. (2018) also observed that the peak amplitude was lower (also indicating a motivational effect) and the peak width was higher in mice prenatally exposed to VPA (prenatal environment) and weaned with other VPA mice (growing environment), particularly in female mice (only at 45 seconds). We did not observe any difference in the peak amplitudes between genotypes. However, consistent with the leftward shift in the timing function of Nrxn1^+/-^, we observed a lower amplitude at 30s for Nrxn1^+/-^ compared to Nrxn1^+/+^ and Nrxn1^ΔS5/-^ mice.

In specific experimental preparations, such as those in which researchers test the acute effects of a drug on timing behavior, the shifts in the timing function are explained in terms of the drug’s effect on the speed of the internal clock. For instance, a leftward shift in the timing function would be attributed to faster clock speed (e.g., Meck, 1996). However, internal clock speed-based explanations do not apply to our findings, since the clock speed by which long-term memory representations of time intervals are established and by which the animal times in the current trial is the same. The clock speed effects would only occur in the case of a mismatch in clock speeds between these two conditions. In fact, the same drug that initially causes a shift in the timing behavior after acute administration would lose its effect after repeated administration since the long-term memory representation would be recalibrated according to the altered clock speed (Meck, 1996).

Shifts in the timing behavior of a genetic model can be explained by the hypothesis of disrupted memory function rather than altered clock speed (see also Gür, Fertan, Alkins, et al., 2019; Gür, Fertan, Kosel, et al., 2019; Karson & Balcı, 2021). For instance, if the consolidation of the memory into the long-term memory is altered such that some of the information is lost (which can be formally exemplified by multiplying the working memory time representation with a value lower than 1 when transferring it to the long-term memory) then the timing behavior would be biased throughout testing. Thus, one explanation for our findings is that Nrxn1^+/-^ mice encode a biased memory representation into long-term memory due to the loss of information (at least compared to Nrxn1^ΔS5/-^ mice). Note that such an account would predict leftward shifts in both start and stop times, which was confirmed by our results in the single-trial analysis. Another possibility is that the long-term memory representation of Nrxn1^+/-^ mice activated with the presentation of a discriminative stimulus leaks over time, which would result in a stronger effect in stop times compared to the start times (as observed in the response curve analysis). Based on the fact that VPA mice did not exhibit a novel object recognition deficit, Acosta et al. (2018) suggested the observed effects in their study were not due to memory deficits in novel object recognition task, however, the task parameters are known to affect the hippocampal involvement in this task (Cohen & Stackman Jr., 2015) and Acosta et al. (2018) did not provide sufficient information regarding the task parameters. Thus, we believe that a biased long-term memory representation remains the most parsimonious explanation of our findings.

Consistent with this explanation, the leftward shift seen in the Nrxn1^+/-^ mice is similar to the effect seen in animals with hippocampal lesions (e.g., Balcı et al., 2009; Olton et al., 1987; Yin & Meck, 2014). Understanding how the hippocampus differs between the brains of ASD individuals and TD individuals is a complex topic. Many studies have examined the volume of the hippocampus in ASD, and the results have been inconsistent. While some have reported increased hippocampal volume (Conti et al., 2020; Groen et al., 2010; Maier et al., 2015; Schumann et al., 2004), others reported no difference (Dager et al., 2007; Nicolson et al., 2006; Piven et al., 1998; van Rooij et al., 2017; Xu et al., 2020), or even a decrease in volume (Aylward et al., 1999; Braden et al., 2017). While a proposed explanation for these diverse findings is that the hippocampus is enlarged in early childhood and normalizes with age (Groen et al., 2010), no difference in hippocampal growth trajectory (Reinhardt et al., 2020), nor significant interaction with age on hippocampal size (Maier et al., 2015; Schumann et al., 2004; van Rooij et al., 2017) have been found.

Several studies have found differences in ASD individuals in the shape of the hippocampus compared to controls (Dager et al., 2007; Nicolson et al., 2006; Richards et al., 2020). Radiomic studies have found differences in the MRI texture of the hippocampus between ASD individuals and controls (Chaddad, Desrosiers, & Toews, 2017; Chaddad, Desrosiers, Hassan, et al., 2017) and decreased density of the hippocampus in ASD (Mei et al., 2020). Post-mortem examinations of ASD hippocampal tissue have found decreased cell size, increased cell body packing, and decreased dendritic branching (Bauman & Kemper, 1985, 2005; Kemper & Bauman, 2002; Raymond et al., 1995). There is also evidence of abnormal neurogenesis and migration (Wegiel et al., 2010) and swollen axon terminals (Weidenheim et al., 2001) in the hippocampi of ASD individuals. Blatt et al.(2001) found decreased density of GABAergic neurons in the hippocampus of ASD brains, while Lawrence et al. (2010) found increased density of GABAergic interneurons.

Studies have found increased connectivity of the hippocampus to the middle temporal gyrus (Rolls et al., 2020), underconnectivity to the perirhinal cortex (Traynor et al., 2018), and reduced whole-brain connectivity during a memory retrieval task (Cooper et al., 2017) in ASD. Increased recruitment of the hippocampus during memory encoding (Hogeveen et al., 2020), and impaired performance on a hippocampal-dependent learning task (Ring et al., 2017) were observed in ASD patients.

Finally, this may be underlain by the increase in the transmission rate (i.e., signal propagation between subsequent time cells) within the time cell architecture that has been found in the hippocampus (Eichenbaum, 2014, 2017). However, this account is inconsistent with the report of decreased excitatory transmission within CA1 pyramidal cells of this ASD model (Lu et al., 2025). Interestingly, the correlational patterns suggested lower relative memory variability or higher relative decision threshold variability in Nrxn1^+/-^ mice.

Another possibility is that altered cortico-striatal circuit function that is highly implicated in interval timing (Buhusi & Meck, 2005), may underlie the timing deficits observed in our work.

Consistent with this view, Acosta et al (2018) reported higher striatal dopamine function and altered dopamine turnover in dorsal striatum (lower in female and higher in male VPA mice) in the VPA model along with leftward shifts in peak response curves similar to our findings. But note that at the information processing level, the related deficits cannot be attributed to the resultant altered clock speed (c.f., Meck, 1996). Testing the same mouse models of ASD in the differential reinforcement of low rates of responding (DRL) task may assist in distinguishing between these possibilities since performance in that task is known to be sensitive to hippocampal dysfunction but relatively less sensitive to striatal dysfunction (e.g., Cho & Jeantet, 2010).

The results gathered in this study (at least the comparison of Nrxn1^-/-^ and Nrxn1^ΔS5/-^ mice) also closely resembled the differences observed between the 5xFAD mice and their wild-type controls (compare Figure 2A in this paper with Figure 2a of Gür, Fertan, Alkins, et al., 2019). This raises the question of whether these similar behavioral effects observed in two very different genetic mouse models of ASD and AD have a common underlying mechanism. In fact, several researchers have focused on the similarities between these two disease states (e.g., Alexiou et al., 2018; Khan et al., 2016; Lahiri et al., 2021; Sokol et al., 2011).

Neurexins also play a role in AD, with levels of NRXN2 and 3 in the CSF proposed as biomarkers of preclinical AD (Lleó et al., 2019), and mutations to NRXN3 have been identified that either increase (Hishimoto et al., 2019), or decrease (Martinez-Mir et al., 2013) susceptibility to AD. Aβ oligomers (AβO) bind to NRXN1 and 2 (Brito-Moreira et al., 2017; Naito et al., 2017), and AβOs reduced NRXN1,2 β expression on axons and inhibited NRXN-mediated synaptic differentiation (Naito et al., 2017). A transgenic mouse model of AD with increased Aβ levels showed decreased expression of NRXN1 and 2 (Naito et al., 2017).

Furthermore, blocking the interaction between AβOs and NRXNs reduced synaptotoxicity induced by AβOs in neuronal cultures and prevented memory impairments in mice injected with AβOs (Brito-Moreira et al., 2017). NRXN1β (Saura et al., 2011) and NRXN3β (Bot et al., 2011) are both processed by α and ɣ secretases, which are important in AD for their role in processing amyloid precursor protein in Aβ. Additionally, mutations associated with familial AD to presenilin 1, a part of the ɣ secretase complex, affect the processing of NRXNs (Bot et al., 2011; Saura et al., 2011).

The fact that a single interval was used in the current study constitutes a limitation of our study. Consequently, we were unable to test whether the core psychophysical features of interval timing, such as the scalar property, are altered in Nrxn1^+/-^ mice. Future studies can test Nrxn1^+/-^ mice in the dual peak interval procedure (as in Acosta et al., 2018). Another limitation of the current study is that only male mice were tested at a single age. Future studies can also test both male and female Nrxn1^+/-^ mice at different ages to better characterize the timing deficit found in Nrxn1^+/-^ mice.

In conclusion, the Nrxn1^+/-^ mice exhibited a leftward shift towards earlier responding on the peak interval task. As the leftward shift is seen in both the start and stop times in the single trial analysis, we believe that a biased long-term memory representation is the most parsimonious explanation for this finding. These results are consistent with findings in mouse and rat VPA experimental (i.e., prenatal exposure) models of ASD and findings in ASD children, providing further evidence of a timing deficit in ASD.

## Contributions

KMR: Formal analysis, Investigation, Data Curation, Writing - Original Draft, Writing - Review & Editing, Visualization; EBH: Investigation, Writing - Review & Editing; REB: Conceptualization, Resources, Writing - Review & Editing, Supervision, Project administration, Funding acquisition; FB: Conceptualization, Methodology, Formal analysis, Resources, Writing - Original Draft, Writing - Review & Editing, Visualization, Supervision, Project administration, Funding acquisition.

## Acknowledgments

We would like to thank Dr Ann Marie Craig of the University of British Columbia for developing and providing the transgenic mice used in this study. Funding was provided by the Natural Sciences and Engineering Research Council of Canada to REB (A7441), the Simons Foundation Autism Research Initiative to REB (608066), and NSERC Discovery Grant (RGPIN-2021-03334) to FB.

## Data availability

Data and analysis code used in this study are available on Borealis, the Canadian Dataverse Repository, at http://doi.org/10.5683/SP3/LWXNZG.

